# Stochastic Differential Equations (SDEs) in NONMEM for Probing Population Pharmacokinetic Model Misspecification: Diagnostic Utility, Practical Considerations, and Future Directions

**DOI:** 10.64898/2026.05.28.728340

**Authors:** Ping Chen, Robert J. Bauer, Yan Li

## Abstract

Population pharmacokinetic (popPK) models are commonly developed using ordinary differential equations (ODEs) to describe deterministic concentration–time profiles, with unexplained variability typically attributed to interindividual variability or residual error. When model misspecification is present, system-level deviations may be absorbed into these conventional variability terms, making the source and magnitude of model inadequacy difficult to assess quantitatively. Stochastic differential equations (SDEs) provide an alternative framework by introducing an explicit system-noise component into the structural model, allowing model–data mismatch to be evaluated more directly. However, historical implementation of SDE-based models in NONMEM has been technically challenging. The availability of the Fortran plug-in subroutine SDE.f90 substantially lowers this barrier and enables more practical implementation of SDE-based models in NONMEM. In this work, SDE-based nonlinear mixed-effects models were evaluated as a quantitative diagnostic framework for probing popPK model misspecification. The SDE.f90 implementation was first verified using simulated one-compartment intravenous bolus datasets with stochastic process noise. Additional simulation–estimation scenarios were then conducted under intentionally misspecified structural or stochastic assumptions, including time-varying elimination, compartmental misspecification, and residual error misspecification. Across these scenarios, the estimated system-noise parameter was generally sensitive to misspecification, with larger values usually associated with greater structural or stochastic mismatch. SDE-based modeling also helped partially separate system-level variability from residual variability and, in selected settings, supported localization of misspecification to specific model components, thereby helping guide model refinement. Overall, SDE-based popPK modeling is a useful addition to the pharmacometric diagnostic toolbox, with system-noise estimates best interpreted together with structural model evaluation, residual diagnostics, parameter behavior, and pharmacologic plausibility.

## Introduction

PopPK modeling plays a central role in characterizing drug exposure and variability in clinical development (1). Traditional popPK models rely on ODEs to describe deterministic concentration-time profiles, with observed data variability attributed to interindividual and residual error variability within a nonlinear mixed-effects framework (2–4). Model misspecification can arise from incorrect structural assumptions such as unmodeled time-varying parameters, inappropriate compartmental representations, misspecified residual error models, and other sources of model–data mismatch (5). In practice, model misspecification is often evaluated indirectly through prior knowledge and experience, diagnostic plots such as visual predictive checks (VPCs), and parameter estimates and associated precision (5, 6). Although these approaches are valuable, they do not provide a direct quantitative framework for identifying or localizing misspecification in a manner that can guide targeted model refinement. Specifically, under the conventional ODE framework, variability arising from model misspecification (system noise) is often absorbed into interindividual or residual error variability, making the source and magnitude of model inadequacy difficult to assess in a statistically explicit manner using standard diagnostics alone.

SDEs provide an alternative modeling framework by explicitly incorporating stochastic system noise into the structural model (7). By introducing a scaling term driven by Brownian motion, SDE-based models allow the concentration-time profile to be described probabilistically instead of deterministically (7, 8). Conceptually, this additional term may capture variability arising from model–data mismatch that is not adequately represented by conventional random-effects components. Specifically, popPK modeling with SDE can introduce an explicit system-noise component (σ_w_∗*dW*) into the PK structural model, thereby creating a potential quantitative indicator of model misspecification. Within a given modeling context, larger estimated system noise may suggest greater mismatch between the assumed model and the underlying data-generating process (9).

The application of SDEs in diagnosing pharmacokinetic model misspecification was demonstrated as early as 2005 by Tornøe et al (10). In that work, SDE-based nonlinear mixed-effects models were used to represent system-level variability arising from structural inadequacies in deterministic models. By introducing system-noise terms into different components of the PK model, the authors showed that the magnitude and location of the system noise could quantitatively indicate both the presence and likely source of model misspecification. For example, a large system-noise estimate associated with the absorption process, combined with negligible estimated system noise in the disposition component, suggested inadequate absorption modeling rather than errors in disposition structure. Subsequent refinement using SDEs further revealed that assumptions such as constant absorption rates were inappropriate, leading to hypothesis-driven model improvements, including time-varying parameters and additional absorption compartments. Once the structural model was improved, the estimated system noise decreased markedly, and parameter estimates became comparable between SDE-and ODE-based models.

Despite this early promising application, broader adoption of SDE approaches remained limited because of substantial implementation complexity in earlier versions of NONMEM (10, 11). At the time, implementing SDE models within NONMEM required manual implementation of Extended Kalman Filter (EKF) algorithms for parameter estimation, and the derivation of covariance matrices for parameters under the SDE framework required extensive additional model-specific mathematical development as well as intensive dataset-preparation steps (7, 10–14). These technical barriers limited practical application to highly specialized users and reduced the accessibility of SDE-based approaches in routine pharmacometric workflows (15, 16).

Advances in NONMEM, beginning with version 7.2 and later versions, now allow direct implementation of SDEs through the Fortran plug-in subroutine SDE.f90, substantially reducing the technical burden that historically restricted their use (12). These developments create an opportunity to revisit the role of SDEs in modern popPK diagnostics. In this work, we examine the value of SDE-based modeling as a quantitative framework for probing popPK model misspecification using the current NONMEM implementation. We first verified the Fortran plug-in subroutine approach for evaluating SDE models, rather than manually implementing the EKF algorithm, then focused on several representative scenarios commonly encountered in popPK modeling, including time-dependent parameter misspecification, compartmental misspecification, and residual error model misspecification. Our goal is not only to highlight the diagnostic potential of SDEs, but also to discuss their practical considerations and the conditions under which interpretation of the estimated system noise should be integrated with broader model evaluation.

## Methods

### Software and Implementation

Data preparation, simulation dataset assembly, and post-processing were performed in RStudio using R version 4.1.3 (Posit Software, PBC, Boston, MA, USA). PopPK ODE and SDE models were implemented in NONMEM® version 7.5.1 (ICON Development Solutions, North Wales, PA, USA). For SDE-based model fitting, the Fortran plug-in subroutine SDE.f90 was used directly in the $SUBROUTINE block, enabling implementation of SDE models without manually deriving and coding extended Kalman filter equations in NONMEM or requiring intensive dataset-preparation steps. Monte Carlo importance sampling expectation-maximization was used for parameter estimation in all NONMEM runs. Representative NONMEM control streams are provided in the Supplementary Codes. Supplementary Code 1 shows the ODE reference model used for comparison in Scenario 1. Supplementary Code 2 shows the one-compartment SDE fitting model with additive system noise applied to the disposition component, which was used as the primary SDE analysis model across the misspecification scenarios. Supplementary Codes 3 and 4 show additional Scenario 1 models used to evaluate localization of system noise to the elimination-rate process.

### Verification of the Fortran Plug-in Subroutine-Based SDE Implementation

To verify the Fortran plug-in subroutine-based SDE implementation in NONMEM, one-compartment IV bolus PK datasets with stochastic process noise were simulated externally in R using the yuima SDE package. The base PK parameters were fixed at CL = 2.3 L/hr and V = 36 L, with an IV bolus dose of 400 mg for 100 subjects. Individual CL and V values were assumed to be log-normally distributed, with an ETA standard deviation of 0.2 for each parameter.

Observed concentrations were generated by applying proportional log-normal residual error with a standard deviation of 0.2.

A proportional SDE process-noise model was used for the amount in the central compartment:

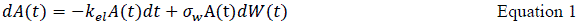

where *A*(*t*) is the amount in the central compartment, σ_*w*_ is the proportional system-noise parameter, and *dW*(*t*) is the Brownian motion increment. In yuima, this was implemented using the drift term-kel*x1 and the diffusion term x1*σ_w_, with the initial amount set to the administered dose. The SDE was simulated over 48 hours using 4800 integration intervals, and concentrations were extracted at nominal sampling times of 0, 0.125, 0.25, 0.5, 1, 1.5, 2, 3, 4, 7, 8, 10, 12, 16, 18, 20, 24, 36, and 48 hours after dose.

The simulated data were then fitted in NONMEM using a one-compartment IV bolus model with the same proportional SDE structure. The Fortran plug-in subroutine SDE.f90 was called directly in the NONMEM $SUBROUTINE block. Parameter recovery was assessed by comparing fitted values with the corresponding simulation truth.

To evaluate the practical recovery range of the proportional SDE parameter, an additional simulation–estimation assessment was conducted in which residual error and interindividual variability were fixed at 0.2, while the true proportional σ_w_ value was varied from 0.05 to 0.25. For the initial verification analysis, parameter recovery was summarized across the tested process noise range, and uncertainty was summarized using standard errors from the NONMEM covariance output when available. The proportional SDE structure was used for the verification analysis, whereas the subsequent misspecification scenarios used additive system noise in the fitted SDE models.

### Common Simulation and Estimation Framework for the Three Scenarios

Three representative simulation–estimation scenarios were conducted under intentionally misspecified structural or stochastic assumptions, including time-varying elimination, compartmental misspecification, and residual error misspecification. Unless otherwise specified, each simulated dataset included 48 subjects, an IV bolus dose of 4000 mg, and nominal sampling times from 0 to 12 hours at 0.5-hour intervals. Individual PK parameters were generated using log-normal interindividual variability, with an ETA standard deviation of 0.2. Observations were generated on the log scale by adding normally distributed residual error, with the residual-error standard deviation varied across simulation settings to evaluate model performance under different noise conditions, including values of 0.1, 0.2, and 0.3.

For each scenario, the simulated data were analyzed using NONMEM models with intentionally misspecified structural or stochastic assumptions. The estimated SDE system-noise parameter, σw, was used to evaluate whether the SDE model could detect model–data mismatch. Parameter estimates and associated uncertainty were extracted from NONMEM output files. Across Scenarios 1–3, different data-generating models were used to create controlled forms of misspecification. However, unless otherwise specified, the primary SDE analysis model used for fitting was the same one-compartment model with additive system noise applied to the disposition component, as shown in Supplementary Code 2. This design allowed the estimated system-noise parameter to be evaluated as a diagnostic signal of mismatch between the fitted one-compartment SDE model and the corresponding data-generating process.

### Scenario 1: Time-Varying Elimination Misspecification

To evaluate whether SDE modeling could detect time-varying elimination misspecification, concentration–time data were simulated using a one-compartment IV bolus model in which the elimination rate constant for each individual decreased exponentially over time. The baseline elimination rate was fixed at 0.25 hr^-1^, and individual elimination rates were generated with log-normal interindividual variability. Time-dependent elimination was introduced as:

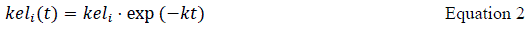

where *k* controlled the severity of time dependence. Ten values of k, ranging from 0 to 0.05, were evaluated. Larger k values produced faster temporal decay in the elimination rate and therefore greater misspecification when the data were fitted using a model assuming constant elimination.

The simulated datasets were fitted using a one-compartment model with a constant elimination rate. Two types of analysis models were evaluated: an ODE model without SDE system noise (Supplementary Code 1) and a one-compartment SDE model with additive system noise applied to the disposition compartment equation (Supplementary Code 2). The ODE model was used to assess how misspecification was absorbed into conventional residual and interindividual variability terms. The SDE model was used to assess whether the estimated σ_w_ increased as the degree of time-varying elimination misspecification increased.

An additional localization analysis was performed by introducing system noise into candidate model components. Three SDE model configurations were evaluated: system noise applied only to the disposition state equation (Supplementary Code 2), system noise applied only to the elimination-rate process (Supplementary Code 3), and system noise applied to both the disposition state equation and the elimination-rate process (Supplementary Code 4). In the model with stochastic elimination, the elimination-rate process was allowed to evolve through an additional diffusion term applied to kel (Supplementary Codes 3 and 4), so that unexplained temporal variation in elimination could be captured by a separate system-noise parameter rather than by concentration-state noise alone. The relative magnitudes of the estimated system-noise terms were compared to assess whether SDE modeling could identify the elimination process as the likely source of misspecification.

### Scenario 2: Compartmental Misspecification

To evaluate compartmental misspecification, datasets were generated from a two-compartment IV bolus model using RxODE. The two-compartment model was defined as:

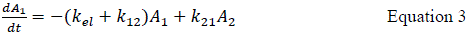

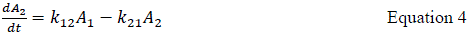

where *A*_1_and *A*_2_ represent the central and peripheral compartment amounts, respectively. The elimination rate *k*_*el*_ was fixed at 0.25 hr^-1^. The inter-compartmental transfer rates *k*_12_and *k*_21_were varied over five values from 0.01 to 1.0 hr^-1^, creating a grid of compartmental-exchange settings. Individual values of *k*_*el*_, *k*_12_, and *k*_21_were generated using log-normal interindividual variability with an ETA standard deviation of 0.2. Proportional log-normal residual error with a standard deviation 0.2 was then added to the simulated central-compartment profiles.

Each two-compartment dataset was then fitted using the misspecified one-compartment SDE model with additive system noise applied to the disposition component (Supplementary Code 2). The purpose was to evaluate whether the estimated σ_*w*_increased as the simulated data became less compatible with a one-compartment structural assumption. The estimated system-noise parameter was summarized across combinations of *k*_12_and *k*_21_to characterize the relationship between compartmental misspecification and SDE-detected system noise.

### Scenario 3: Residual Error Model Misspecification

Residual error model misspecification was evaluated using one-compartment IV bolus datasets generated under three alternative residual error structures and fitted using an analysis model assuming independent proportional residual error. The base structural model used a typical elimination rate of 0.25 hr^-1^, log-normal interindividual variability with an ETA standard deviation 0.2, and an IV bolus dose of 4000 mg.

First, combined residual error misspecification was evaluated by generating observations with proportional log-normal residual error and an additional additive error contribution. The proportional residual error standard deviation was fixed at 0.3, and the additive error contribution was varied across increasing values. The simulated data were then fitted using the one-compartment SDE model with additive system noise and proportional residual error (Supplementary Code 2).

Second, additive residual error misspecification was evaluated by generating data with an additive error contribution alone, without proportional residual error. The additive error magnitude varied across increasing values, and all datasets were fitted using the same one-compartment SDE model with additive system noise and proportional residual error (Supplementary Code 2). This created a controlled mismatch between the true additive residual error structure and the fitted proportional residual error assumption.

Third, autocorrelated residual error misspecification was evaluated by generating residual errors with serial correlation and fitting the data using an independent proportional residual error model. Autocorrelated residual errors were generated using multivariate normal residual vectors with covariance matrices determined by the autocorrelation parameter ρ, which varied from 0 to 0.99. AR(1), AR(2), and AR(3)-type covariance structures were evaluated. For AR(1), the covariance between residuals decreased as a function of ρ^∣*i*−*j*∣^. For AR(2) and AR(3)-type structures, additional lagged correlation components were included, and the covariance matrix was adjusted to be positive definite before simulation. The resulting datasets were analyzed using the same one-compartment SDE model with additive system noise and independent proportional residual error (Supplementary Code 2).

Across the residual error misspecification settings, the estimated σ_*w*_was summarized to evaluate whether SDE system noise increased as the discrepancy between the true and assumed residual error structures increased.

### Assessment of Limitations of SDE-Based Diagnostics

Additional simulations were conducted to identify conditions under which the estimated SDE system-noise term became less directly interpretable. These analyses focused on two broad limitation settings: extreme model misspecification and high variability conditions.

For extreme structural misspecification, the compartmental misspecification scenario was examined across increasingly divergent two-compartment parameter settings. The purpose was to evaluate whether the estimated σ_*w*_continued to increase with worsening structural mismatch or whether misspecification-related variability was redistributed into other model components, such as fixed-effect elimination or conventional interindividual variability.

For extreme residual error misspecification, the autocorrelated residual error scenario was examined at high autocorrelation values. Datasets generated with strongly autocorrelated residual errors were fitted using models assuming independent residual error. Estimated σ_*w*_, residual error, and interindividual variability were examined to assess whether the SDE system-noise term remained a reliable indicator of residual structure misspecification under severe autocorrelation.

## Results

### Verification of the Fortran Plug-in Subroutine-Based SDE Implementation in NONMEM Using Simulated One-Compartment IV PK Data

Because published NONMEM applications of SDE models have historically relied largely on manual EKF implementation and extensive data preprocessing, an initial simulation–estimation analysis was conducted to evaluate whether the Fortran plug-in subroutine SDE.f90 could be used directly in NONMEM to recover externally generated stochastic process variability from a one-compartment IV bolus PK model. This approach was evaluated as a practical alternative to manually deriving and coding extended Kalman filter equations in NONMEM, which is technically challenging and difficult to generalize.

Representative simulated profiles were generated in R using the yuima package and shown in Supplementary Figure 1A. For six randomly selected subjects, deterministic ODE profiles without stochastic process noise or residual error were overlaid with profiles including proportional SDE process noise (proportional system noise, σw = 0.2) and profiles including both SDE process noise and residual error (proportional residual error, σ = 0.2). The simulated profiles showed the expected pattern: the SDE component introduced trajectory-level deviations from the deterministic ODE profile, whereas residual error introduced additional observation-level variability around the stochastic trajectory.

Parameter recovery was then evaluated under fixed moderate interindividual variability and residual error, both set to 0.2 on the log scale, while the true proportional SDE parameter, σ_w_, varied from 0.05 to 0.25. As shown in Supplementary Figure 1B, estimated σ_w_ values were generally aligned with the true simulated σ_w_ values, although slight underestimation was observed at the higher end of the evaluated range. Consistent with this pattern, estimated residual error (σ) was well recovered, although it became slightly overestimated at higher σ_w_ values (Supplementary Figure 1C). Similarly, the estimated interindividual variability, particularly for clearance, increased at higher σ_w_ values (Supplementary Figure 1D), indicating increasing confounding among stochastic process noise, residual variability, and interindividual variability.

Overall, these results support the feasibility of using the Fortran plug-in SDE.f90 routine in NONMEM for proportional SDE-based PK modeling. Reasonable recovery was observed across the tested range of process noise, whereas higher process-noise levels led to slight underestimation of σ_w_ and compensatory inflation of residual error and interindividual variability, particularly for clearance. These findings provide initial verification that direct implementation of the Fortran plug-in SDE.f90 routine performs as expected within a practically identifiable range and supports further simulation-based evaluation under broader model and study-design settings.

### Scenario 1: Detection and Localization of Time-Varying Elimination Misspecification

The first simulation–estimation scenario was conducted under intentional structural misspecification, in which data generated with time-varying elimination were fitted using models assuming a constant elimination rate. Specifically, concentration–time data were generated from a one-compartment model in which the elimination rate constant, kel, decreased exponentially over time, whereas the analysis model assumed a constant kel. This created a controlled form of structural misspecification whose severity increased as the decay parameter, k, increased. As shown in Figure 1A, larger k values produced progressively faster decay of over time; for example, when k = 0.045 hr⁻¹, kel at 48 hours was approximately 10% of its initial value at time 0. This increasing deviation from the constant-elimination assumption represented progressively greater structural misspecification.

**Figure 1.**
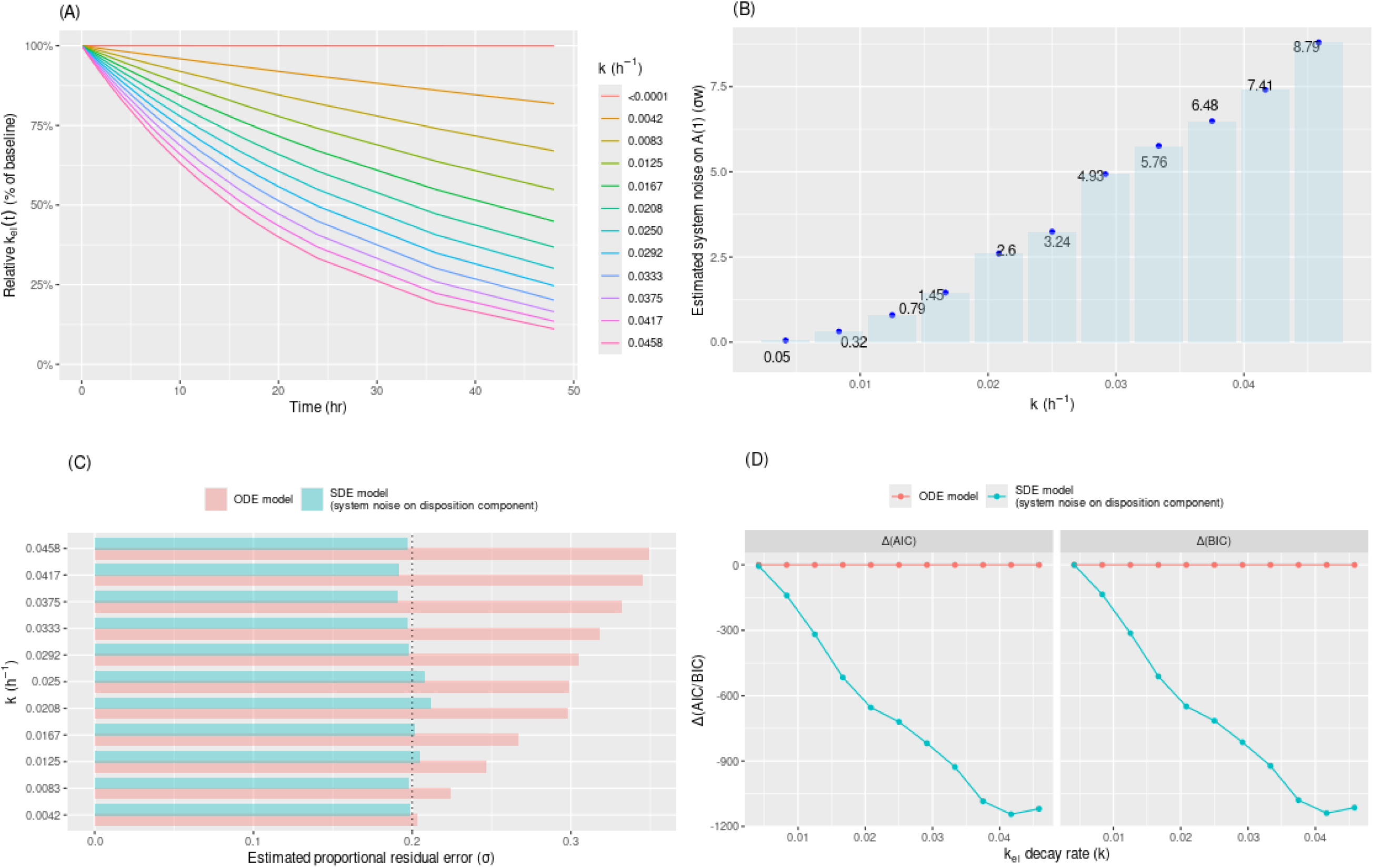
Detection of time-varying elimination misspecification using SDE-based models in Scenario 1. (A) Simulated time-varying elimination-rate profiles under different values of the decay parameter, k. The y-axis shows the elimination rate as a percentage of its initial value at time 0. Larger k values indicate faster temporal decay in kel and therefore greater departure from the constant-elimination assumption. (B) Estimated additive system noise, σw, from SDE-based models with system noise applied to the disposition component. Increasing σw values indicate increasing model–data mismatch as the degree of time-varying elimination misspecification increases. (C) Estimated proportional residual error (σ), from ODE– and SDE-based models across different k values. The dashed vertical reference line indicates the true residual error value used in simulation. Comparison of the ODE and SDE models illustrates whether misspecification-related variability is absorbed into residual error or partially captured by the SDE system-noise term. (D) Difference in information criteria between SDE– and ODE-based models across different k values. ΔAIC and ΔBIC were calculated as the information criterion value for the SDE model minus that for the ODE model. More negative values indicate stronger support for the SDE-based model relative to the ODE-based model.

When these datasets were analyzed using SDE-based models with system noise applied to the disposition component, the estimated system-noise term, σ_w_, increased systematically with increasing k (Figure 1B). In contrast, when no time dependency (k = 0) was introduced in the elimination process, the estimated system noise remained close to zero, consistent with an adequately specified model. These findings indicate that, within this scenario, the SDE framework was able to quantitatively detect the presence of time-varying elimination misspecification, and that the magnitude of the estimated system noise tracked the severity of the model–data mismatch.

In addition to detecting structural model misspecification and providing a quantitative measure of its magnitude, the SDE framework facilitated clearer partial separation of system noise from conventional residual variability. Under the corresponding ODE-based model fits, the estimated residual error variability increased as the degree of time-varying elimination misspecification increased (pink bars in Figure 1C), indicating that misspecification-related variability was being absorbed into the residual variability term. By comparison, the SDE-based model fits maintained residual error estimates closer to the true value across increasing levels of misspecification (light blue bars in Figure 1C), while assigning part of the unexplained variability to the explicit system-noise term. Consistently, both ΔAIC and ΔBIC comparing the SDE and ODE models decreased substantially as the degree of misspecification increased (Figure 1D). These information criteria were used as supportive summaries under the implemented likelihood framework and suggested improved accommodation of model–data mismatch after adding the system-noise term; however, the primary diagnostic interpretation was based on the coordinated behavior of σw, residual error, parameter estimates, and visual model behavior rather than information criteria alone.

SDE-based modeling also improved the description of individual concentration–time profiles under the misspecified constant-elimination model. As illustrated in Figure 2, individual predictions from the ODE-based model followed deterministic profiles, whereas predictions from the SDE-based model were able to adapt more closely to the observed data. This behavior reflects the conditional nature of the SDE-based prediction framework, in which the latent trajectory can be informed by observed data through the filtering process instead of being constrained to a fully deterministic ODE trajectory.

**Figure 2.**
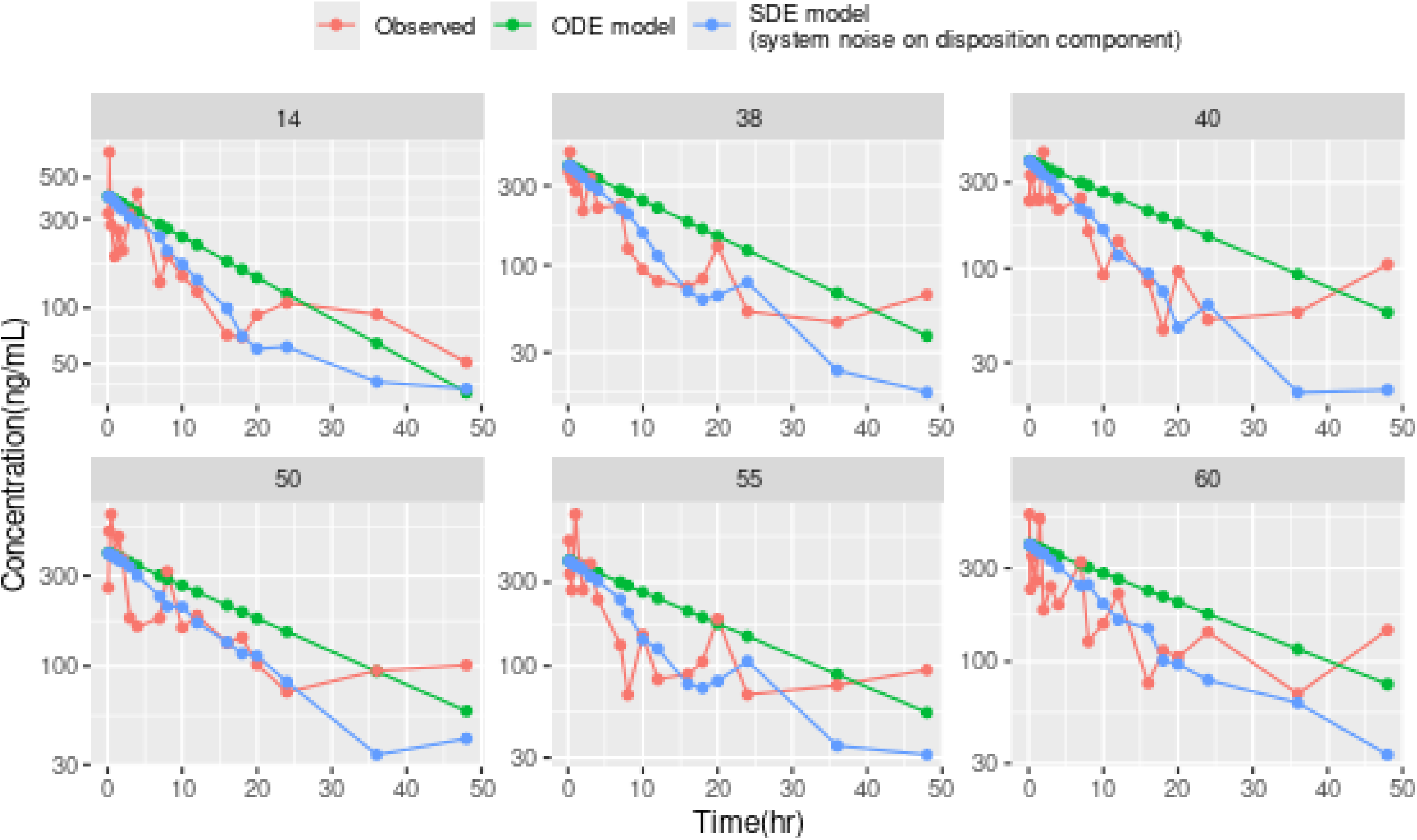
Representative individual concentration–time profiles comparing ODE– and SDE-based models in Scenario 1. Six representative individual profiles are shown for Scenario 1 under k = 0.05 hr⁻¹ and residual error σ = 0.2. Observed concentrations are shown as points, ODE-based individual predictions are shown in green, and SDE-based individual predictions are shown in blue.

To further evaluate whether the SDE approach could help localize the source of misspecification, system noise was introduced into different model components. When system noise was applied only to the disposition state equation, the estimated disposition-related system noise was large and increased with k, indicating substantial unexplained model–data mismatch (Figure 1B and blue bars in Figure 3A). However, when an additional system-noise term was introduced on the elimination-rate process, the system-noise estimate associated with the disposition component decreased markedly to a negligible level (red bars in Figure 3A), while the system-noise estimate associated with the elimination process increased with k (Figure 3B). Posterior estimates of kel further showed clear temporal variation in the elimination rate constant (Figure 3C). In this controlled simulation setting, this pattern correctly recovered the imposed time-varying elimination mechanism and supported that the dominant mismatch arose from the constant-elimination assumption rather than from the disposition state equation itself.

**Figure 3.**
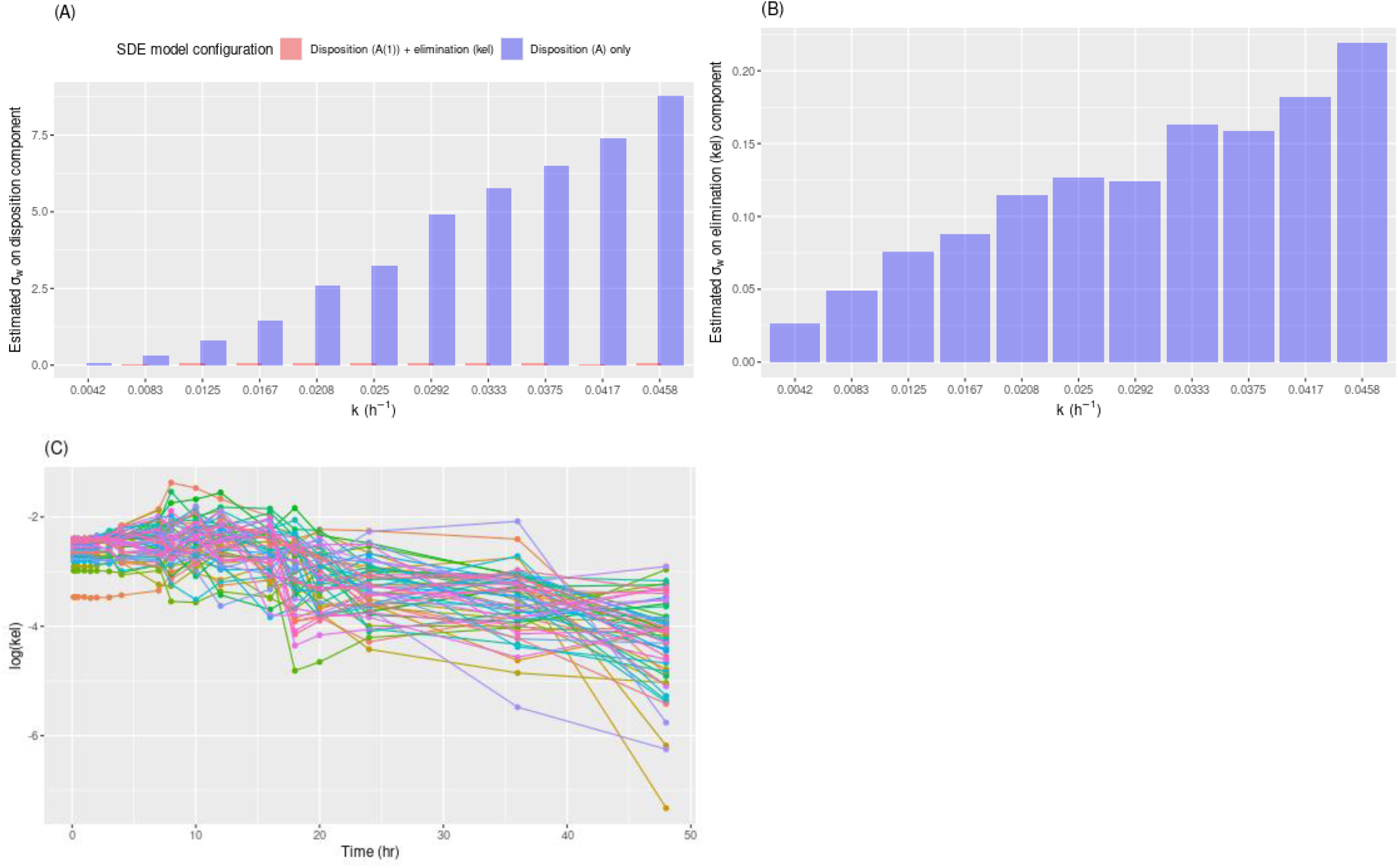
Localization of time-varying elimination misspecification using component-specific SDE system noise. (A) Estimated system noise on the disposition component across different k values. Results are shown for models with system noise applied to the disposition component only and for models with system noise applied to both the disposition component and the elimination-rate process. The marked reduction in disposition-related system noise after adding system noise to the elimination-rate process suggests that the primary mismatch is not localized to the disposition state equation itself. (B) Estimated system noise on the elimination-rate process across different k values. Increasing estimates indicate that unexplained temporal variation is primarily captured by the stochastic elimination-rate component. (C) Posterior estimates of log(kel) over time from the SDE-based model with system noise applied to the elimination-rate process. The temporal pattern is consistent with the known time-varying elimination mechanism used in the simulation and supports localization of the imposed misspecification to the constant-elimination assumption.

Taken together, this scenario shows that SDE-based modeling was able not only to detect the presence of time-varying elimination misspecification, but also to provide mechanistic insight into its likely origin. In this setting, the estimated system noise served as both a quantitative indicator of model inadequacy and could provide a useful guide for hypothesis-driven model refinement.

### Scenario 2: Detection of Compartmental Misspecification

In the second scenario, concentration–time data were simulated from a two-compartment model and subsequently analyzed using a misspecified one-compartment model augmented with system noise. The degree of apparent compartmental misspecification was explored by varying combinations of the inter-compartmental exchange parameters k_12_ and k_21_, which generated profiles with different degrees of distribution-phase behavior and different levels of compatibility with a one-compartment approximation. As shown in Figure 4A, small k₂₁ values, particularly when combined with larger k₁₂ values, produced more pronounced distribution-phase behavior and greater deviation from a one-compartment profile. These profiles could not be adequately captured by a one-compartment structural model.

**Figure 4.**
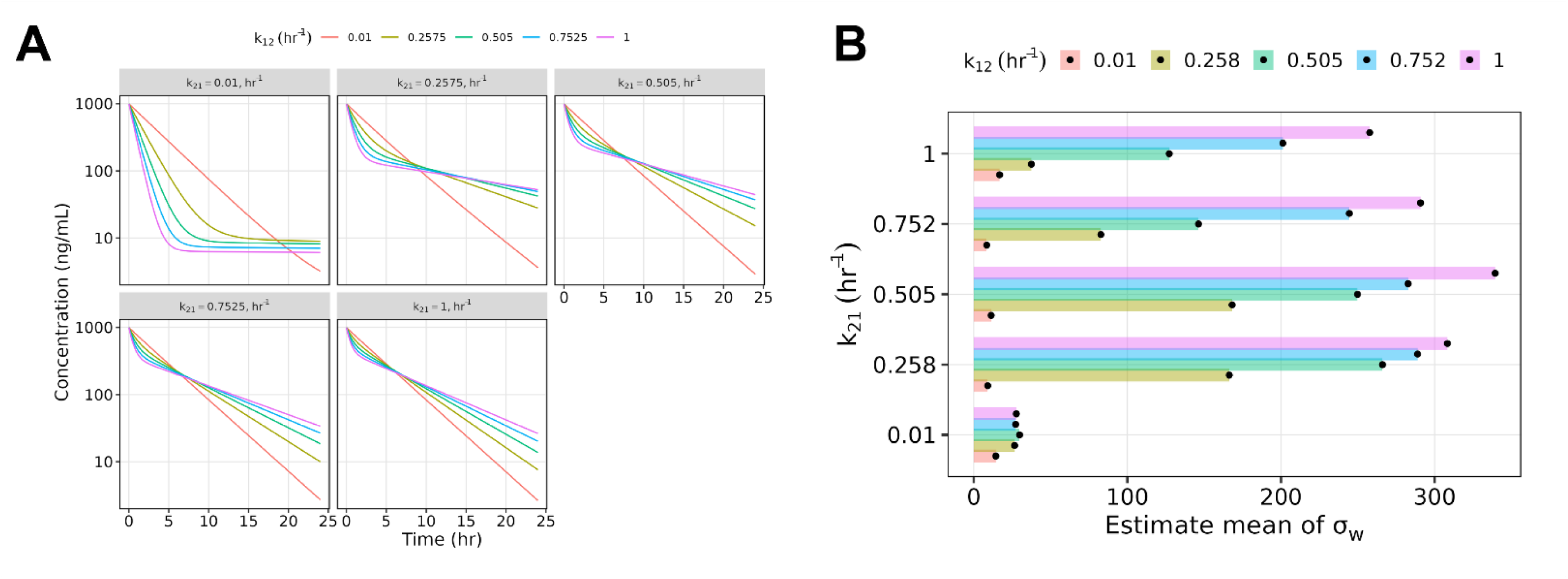
Detection of compartmental misspecification using one-compartment SDE-based models in Scenario 2. (A) Simulated concentration–time profiles generated from two-compartment IV bolus models under different combinations of k₁₂ and k₂₁. Colors represent different k₁₂ values, and panels represent different k₂₁ values. Profiles with more pronounced distribution-phase behavior indicate greater deviation from a one-compartment structural assumption. (B) Estimated system noise, σw, from misspecified one-compartment SDE-based models fitted to the two-compartment simulated datasets. Colors represent different k₁₂ values, and the y-axis represents k₂₁ values. Larger σw estimates indicate greater model–data mismatch captured by the SDE system-noise term.

When fitted with an SDE-based one-compartment model, the estimated system noise (σ_w_) generally increased as the simulated profiles became more distinctly two-compartment in nature (Figure 4B). The largest σ_w_ estimates were observed in settings with more pronounced compartmental-exchange imbalance, consistent with greater structural mismatch between the two-compartment data-generating model and the one-compartment analysis model. These findings indicate that the SDE framework was sensitive to compartmental misspecification and that the estimated system noise broadly reflected the severity of model–data mismatch.

However, the relationship between compartmental misspecification and estimated σ_w_ was not strictly monotonic across all combinations of k₁₂ and k₂₁. In extreme cases, particularly when the simulated one-compartment approximation was very far from the true two-compartment behavior, the estimated system noise was lower than would have been expected based on the degree of structural mismatch. This suggests that, under more severe compartmental misspecification, part of the mismatch may be absorbed by other model components, including fixed-effect parameter estimates or conventional random-effects terms.

Taken together, these findings indicate that SDE-based modeling can detect compartmental misspecification and provide a quantitative signal of structural mismatch under a broad range of conditions. At the same time, the estimated system-noise parameter should be interpreted together with the overall model behavior, because σw may not serve as a universally monotonic measure of misspecification severity when the analysis model is very far from the true data-generating structure.

### Scenario 3: Sensitivity of SDE System Noise to Residual Error Model Misspecification

In the third scenario, we evaluated whether the estimated SDE system-noise parameter was sensitive to residual error model misspecification under three settings: combined residual error fitted with a proportional error model, additive residual error fitted with a proportional error model, and autocorrelated residual error fitted with an independent residual error model. In all cases, the analysis model included an explicit system-noise term to assess whether misspecification in the residual structure could be reflected by the estimated system noise.

When data generated under a combined residual error model were analyzed using a proportional error model, the estimated system-noise term increased as the additive error contribution increased (Figure 5A). This pattern indicates that the estimated σ_w_ was sensitive to variability introduced by the unmodeled additive residual-error component. Similarly, when data generated under a purely additive residual error model were fitted using a proportional error model, the estimated system noise also increased with increasing incompatibility between the true and assumed residual error structures (Figure 5B). Taken together, these results suggest that the system-noise term can absorb part of the unexplained variability introduced by an incorrect residual error specification and thereby serve as a useful indicator of unmodeled stochastic structure or residual-model misspecification.

**Figure 5.**
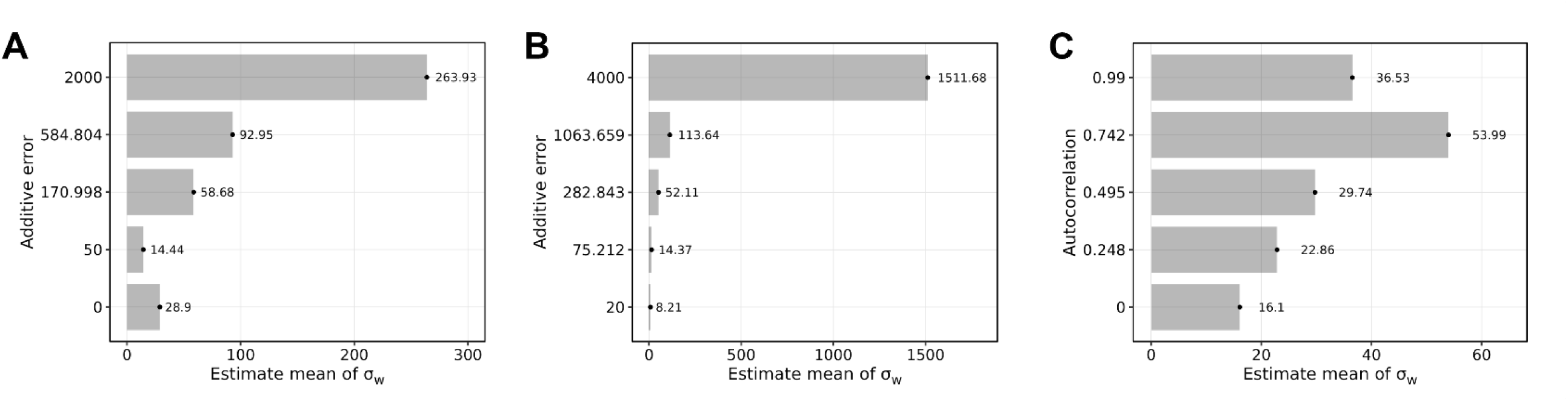
Sensitivity of SDE system-noise estimates to residual error model misspecification in Scenario 3. (A) Estimated system noise, σw, for data generated using a combined residual error model and fitted using a proportional residual error model. (B) Estimated system noise, σw, for data generated using an additive residual error model and fitted using a proportional residual error model. (C) Estimated system noise, σw, for data generated with autocorrelated residual errors and fitted using an independent residual error model. Across panels, larger σw estimates indicate that the SDE system-noise term captured part of the unexplained variability introduced by residual error model misspecification.

A similar pattern was observed when the true residual errors were autocorrelated, but the analysis model assumed independent residual variability. As the autocorrelation in the simulated data increased, the estimated system-noise term generally increased as well (Figure 5C), indicating that the SDE-based model was sensitive to unmodeled serial dependence in the residual structure. This finding is relevant because residual autocorrelation is a common indication of inadequacy in the assumed stochastic model, yet it may not be readily quantified within a conventional ODE-based framework. Although the relationship was not strictly monotonic at the highest autocorrelation level, the overall trend supported sensitivity of the estimated system-noise term to residual autocorrelation.

Across the residual error misspecification settings, SDE-based modeling demonstrated sensitivity to discrepancies between the true and assumed residual error structures. However, an increased σ_w_ in this setting should not be interpreted as direct evidence of true biological or process noise. Rather, it indicates that the SDE term can capture part of the unexplained variability induced by an incorrect residual-error assumption. Therefore, in residual error misspecification scenarios, estimated system noise should be interpreted as a diagnostic signal of unmodeled stochastic structure instead of as definitive evidence of underlying physiological process variability.

### Limitations of SDE-Based Diagnostics

Although the preceding scenarios support the utility of SDE-based modeling for detecting model misspecification, additional simulations identified settings in which the estimated system-noise term may no longer scale consistently with misspecification severity or remain directly interpretable. Two broad boundary conditions emerged: substantial model misspecification and high-variability data conditions. Several of these scenarios were intentionally designed as extreme stress tests, with the goal of challenging the limits of the current SDE implementation and evaluating when the diagnostic signal may begin to deteriorate. In both cases, misspecification-related variability was not captured primarily by the explicit system-noise term but instead was partly redistributed into other model components.

Under substantial structural misspecification, the estimated system noise could become lower than expected despite clear model inadequacy. In the compartmental misspecification setting, when the one-compartment analysis model was substantially different from the true two-compartment data-generating model, the mismatch was partially absorbed into the fixed-effect elimination parameter rather than continuing to accumulate in the system-noise term (Figure 6A–B). As a result, the estimated system noise no longer increased in proportion to the severity of the structural discrepancy. A similar phenomenon was observed in the residual error setting. When the true residual process was highly autocorrelated, but the fitted model assumed independent residual variability, the misspecification was partially absorbed into increased interindividual variability, again leading to lower-than-expected system-noise estimates under the most extreme conditions (Figure 6C–D). These results should therefore be interpreted as boundary-case behavior under intentionally exaggerated misspecification, instead of as evidence that SDE diagnostics are unreliable under ordinary modeling conditions.

**Figure 6.**
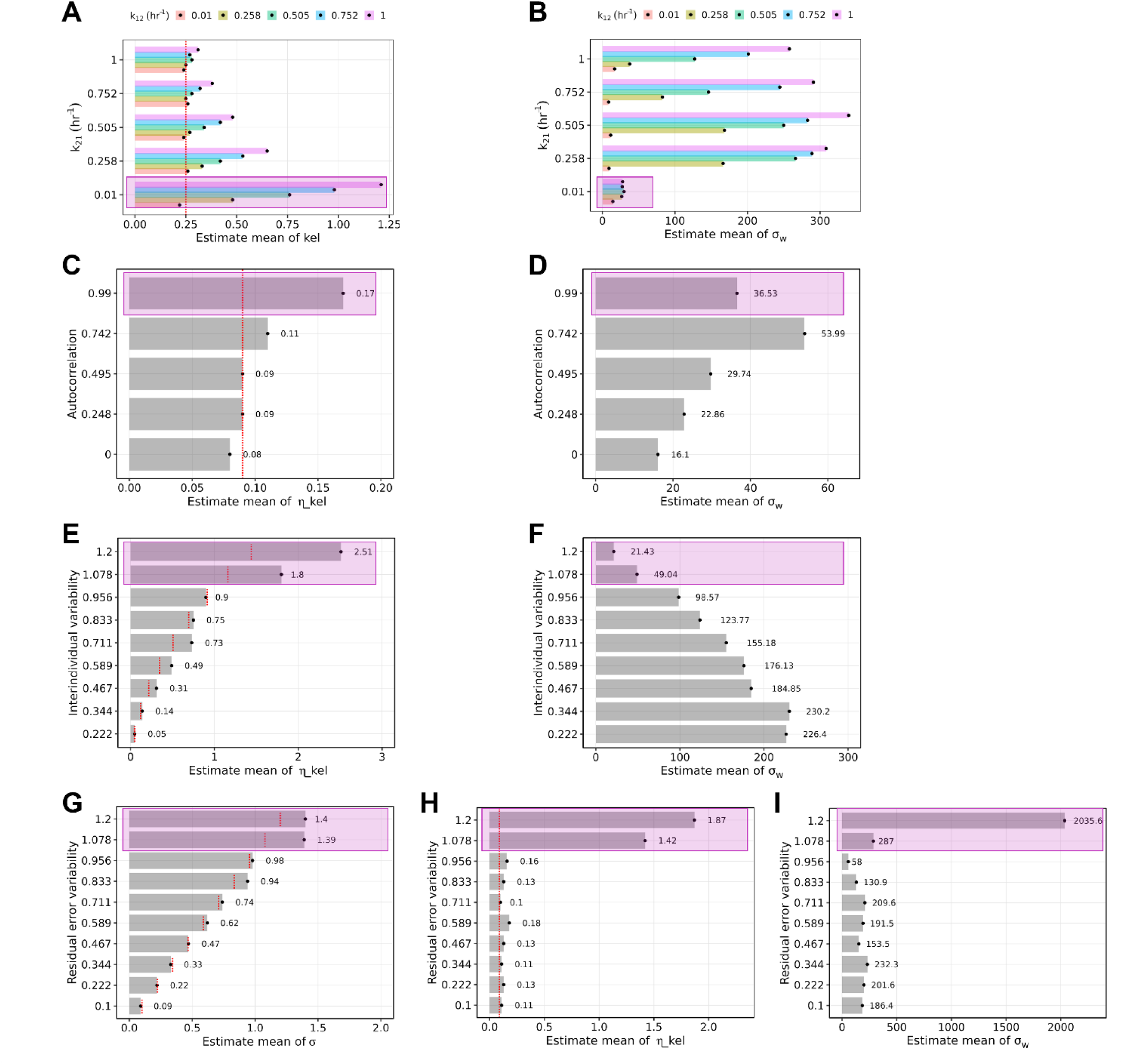
Boundary conditions affecting interpretability of SDE system-noise estimates. Purple shaded regions indicate extreme settings in which estimated system noise no longer increased proportionally with worsening misspecification or increasing variability. Panels A–B show compartmental misspecification; panels C–D show autocorrelated residual error misspecification; panels E–F show high interindividual variability; and panels G–I show high residual error variability. Red dashed lines indicate true simulated values where applicable. Panel-specific outputs include estimated elimination rate constant (A), system noise (B, D, F, I), interindividual variability (C, E, H), and residual error variability (G).

High-variability data conditions also reduced the interpretability of the system-noise estimate. When interindividual variability was made extremely large, variability arising from model misspecification became increasingly masked by the random-effects structure, such that the estimated system noise no longer rose as expected with worsening misspecification (Figure 6E–F). Likewise, when residual error variability became extremely high, model fitting became less stable and the separation among interindividual variability, residual error variability, and system noise became less distinct (Figure 6G–I). Under these conditions, the estimated system noise was influenced not only by model misspecification but also by the reduced ability of the model to decompose total variability into interpretable components. As with the structural misspecification examples, these high-variability simulations were intended to pressure-test the framework by creating conditions in which identifiability and variance decomposition would be expected to become challenging.

Taken together, these findings indicate that the diagnostic value of the SDE framework is context dependent instead of absolute. Under moderate and interpretable misspecification, the system-noise term behaved as an informative indicator of model inadequacy. However, when the assumed model became too far removed from the true system, or when data variability became excessively large, misspecification-related signals were redistributed across fixed effects and conventional random-effects terms, thereby weakening the direct interpretability of the estimated system noise.

## Discussion

The present study evaluated SDE-based popPK modeling in NONMEM as a quantitative diagnostic framework for probing model misspecification that may otherwise be difficult to identify or localize using conventional ODE-based approaches. In traditional popPK models, concentration–time profiles are described deterministically, and unexplained variability is typically attributed to interindividual variability or residual error (2–4). When misspecification arises from incorrect structural assumptions, standard diagnostics may indicate model inadequacy but may not clearly identify its source or provide a quantitative basis for targeted model refinement (5, 6). SDE-based modeling has been shown in earlier work to provide a useful framework for this purpose by introducing an explicit system-noise term into the structural model. However, despite these promising early applications, broader adoption of SDE-based modeling remained limited because earlier NONMEM workflows required manual EKF implementation, modified data structures, and specialized mathematical derivations (10). The current study is therefore timely because modern NONMEM implementations with the Fortran plug-in subroutine SDE.f90 have substantially reduced this technical barrier, making broader evaluation of SDE-based approaches more practical. Nevertheless, greater implementation convenience does not eliminate the need to understand the diagnostic behavior and practical limitations of estimated system-noise terms (14, 17, 18).

In the present study, the initial simulation–estimation verification showed that the NONMEM Fortran plug-in routine SDE.f90 could recover SDE process noise with reasonable accuracy over a practically identifiable range. This finding supports the feasibility of direct SDE implementation in NONMEM using the Fortran plug-in subroutine. Across the evaluated scenarios, the estimated system-noise term generally increased when the fitted model became less consistent with the data-generating process and remained minimal when the model assumptions were aligned with the simulated data. This behavior supports the diagnostic rationale for SDE-based modeling, in which part of the model–data mismatch can be represented as system-level uncertainty within the state dynamics instead of being absorbed entirely into interindividual or residual variability.

A key strength of this work is that it extends prior SDE applications into a systematic evaluation of multiple misspecification scenarios using the current NONMEM SDE.f90 implementation. Earlier studies, including the 2005 degarelix analysis, showed that SDEs could identify structural inadequacy in absorption, track temporal changes in absorption behavior, and guide refinement from a simple first-order absorption model to a more appropriate multi-component absorption model (10). In that study, the relevant diffusion terms decreased after structural model refinement, illustrating that SDEs can serve as a model-development tool when interpreted in the context of model structure and pharmacologic plausibility. The present results are consistent with that diagnostic logic and broaden its application beyond absorption-related misspecification to time-varying elimination, compartmental misspecification, and residual error misspecification. Importantly, the value of SDE modeling was not limited to estimating a single system-noise parameter. In Scenario 1, the framework did more than detect misspecification: it helped localize the source of the mismatch to the elimination process. This is important because one of the central limitations of conventional ODE-based fitting is that misspecification-related variation is typically redistributed into conventional variability terms, making it difficult to distinguish model inadequacy from biological heterogeneity or measurement noise (19). The current findings therefore support the view that, when used thoughtfully, SDEs can provide mechanistic guidance for model refinement (9).

Our findings are also consistent with prior PK/PD applications in which the practical benefits of SDE extended beyond model fit (20, 21). In the IL-21 thermoregulation work, SDEs improved the autocorrelation structure of residuals, produced more realistic simulations than ODE-based models, and in some cases reduced the need to invoke separate inter-occasion variability terms because the system-noise process captured continuous day-to-day fluctuation more naturally (13). Likewise, in the OGTT analysis, adding an OU-type stochastic component reduced autocorrelation of one-step prediction errors and improved the relationship between model-derived beta-cell function indices and IVGTT-derived AIR0–8, including their relationships with covariates (22, 23). These studies suggest that the benefit of SDE is not merely descriptive; under appropriate conditions, it can improve the scientific interpretability and predictive relevance of the fitted model.

Three points are worth emphasizing regarding the interpretation of SDE-based diagnostics. First, the present work shows that the estimated system-noise term should not be interpreted as an absolute or universally monotonic measure of misspecification severity. In several extreme settings, worsening misspecification did not lead to correspondingly larger system-noise estimates. Instead, the mismatch was partially absorbed into other model components, including fixed-effect parameters, interindividual variability, and residual error. This behavior is expected from an estimation perspective: when the assumed model is far from the true data-generating process, multiple model components may compete to absorb the same discrepancy. Thus, SDE diagnostics appear most informative when the candidate model remains sufficiently close to the underlying system for the estimated system noise to retain mechanistic interpretability. Second, SDE-based diagnostics are sensitive to **data informativeness and the relative magnitude of variability components**. The current simulations showed that very large residual variability can mask the contribution of model misspecification to the system-noise term. **Similarly, large interindividual variability can reduce the ability to distinguish systematic model–data mismatch from subject-level heterogeneity.** When a large portion of the observed variability is already captured by broad random-effects terms or high residual noise, the model has less ability to isolate system-level mismatch as a separate component. This limitation is practically important because SDE-based diagnostics require sufficient information to decompose total variability into interpretable sources. Accordingly, SDE diagnostics may be most useful in data settings with adequate signal-to-noise ratio, where system-level mismatch can be distinguished from interindividual variability and observation-level error (24). Third, SDE-based modeling is not a substitute for mechanistic thinking. A large system-noise estimate may indicate that the fitted model is inadequate, but it does not by itself explain why. The strength of the approach emerges when system noise is introduced strategically into candidate model components and interpreted alongside pharmacology, physiology, and model structure. The degarelix absorption example illustrates this point: the key finding was not simply that σw was large, but that the pattern of system-noise estimates across model components suggested a time-varying absorption process and motivated a more physiologically plausible structural refinement. Similarly, the gastric-fluid-volume analysis used time-varying stochasticity on gastric emptying as a diagnostic strategy to identify the main driver of system dynamics. Therefore, SDE-based modeling is most useful when used to generate and test mechanistic hypotheses, not as a purely empirical compensation mechanism for an inadequate structural model.

From a pharmacometric workflow perspective, the present results support a practical role for SDE-based diagnostics as a selective, complementary tool (1, 25). They are probably not needed for every routine popPK analysis, particularly when standard structural assumptions are adequate and residual diagnostics are well behaved. Instead, SDE-based diagnostics are likely most useful when standard structural assumptions remain uncertain, residual diagnostics show persistent serial structure, time-varying physiology is plausible, compartmental behavior is not well captured, or conventional variability terms appear to be absorbing systematic model–data mismatch (26, 27). In these settings, SDE models can provide an additional diagnostic layer to help distinguish random variability from systematic model inadequacy and support hypothesis-driven model refinement. Thus, SDE-based modeling may be best positioned as an escalation tool when conventional model development and diagnostics do not fully explain the observed data patterns.

This work also points to several future directions. The availability of SDE.f90 in NONMEM substantially lowers the implementation barrier and makes SDE-based modeling more accessible alongside established diagnostics, including goodness-of-fit plots, VPCs, residual trends, parameter precision, and sensitivity analyses. However, broadly accepted operating criteria for interpreting the magnitude or significance of the system-noise term are still lacking. Future work should therefore focus on scenario-dependent calibration, simulation-based reference ranges, empirical thresholds, and comparative evaluation across competing models, rather than relying on absolute σ_w_ cutoffs. Translation to real clinical datasets will also be essential to determine whether SDE-based diagnostics can improve recognition of structural inadequacy and guide more efficient model refinement in practice. Particular opportunities may exist in highly dynamic systems, including oral absorption, gastrointestinal transit, time-varying clearance, disease progression, cell-therapy kinetics, and other biological settings where deterministic ODE assumptions may be strained (28). More broadly, future work should evaluate when SDE-based diagnostics provide incremental value beyond standard diagnostics and how their outputs should be interpreted in clinical pharmacology and regulatory contexts (11).

## Conclusions

SDE-based popPK models provide a useful quantitative approach for detecting and, in some settings, localizing model misspecification, thereby helping make system-level model–data mismatch more visible and partially separable from conventional interindividual and residual variability. By explicitly modeling system noise, SDEs provide diagnostic information that is not directly available from conventional ODE-based diagnostics alone under suitable conditions. The availability of the Fortran plug-in subroutine SDE.f90 has substantially lowered the technical barrier for implementing SDE-based models in NONMEM, making this approach more accessible for practical pharmacometric applications. Overall, these findings support SDE-based modeling as a practical addition to the broader popPK diagnostic framework, while recognizing that estimated system noise should be interpreted together with structural model evaluation, residual diagnostics, parameter behavior, and pharmacologic plausibility.

## Notes

### Competing Interest Statement

The authors have declared no competing interest.

### Funding

- Bristol-Myers Squibb (United States)

