## Supplementary for "Stochastic Differential Equations (SDEs) in NONMEM for Probing Population Pharmacokinetic Model Misspecification: Diagnostic Utility, Practical Considerations, and Future Directions"

**Figure Legends**

**Supplementary Figure 1: Verification of NONMEM SDE.f90 implementation using externally simulated one-compartment IV bolus SDE data.** (A) Representative simulated concentration–time profiles generated in R using the yuima package. Deterministic ODE profiles without stochastic process noise or residual error are overlaid with profiles including proportional SDE process noise and profiles including both proportional SDE process noise and residual error. (B) Recovery of the proportional system-noise parameter, comparing true simulated σ_w_ with NONMEM-estimated σ_w_. The dotted reference line represents the line of identity. (C) Recovery of the proportional residual error σ across increasing true simulated σ_w_ values. The dotted horizontal reference line represents the true residual error σ used in simulation. (D) Recovery of interindividual variability parameters for clearance and volume across increasing true simulated σ_w_ values. The dotted horizontal reference lines represent the true ETA variance values used in simulation.

**Supplementary Figure 1: Verification of NONMEM SDE.f90 implementation using externally simulated one-compartment IV bolus SDE data.**

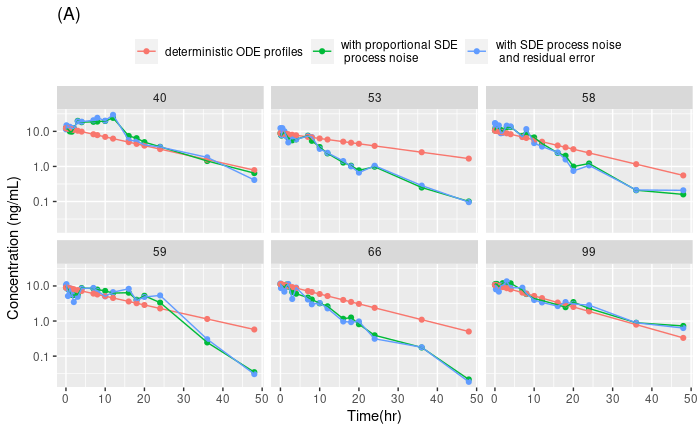

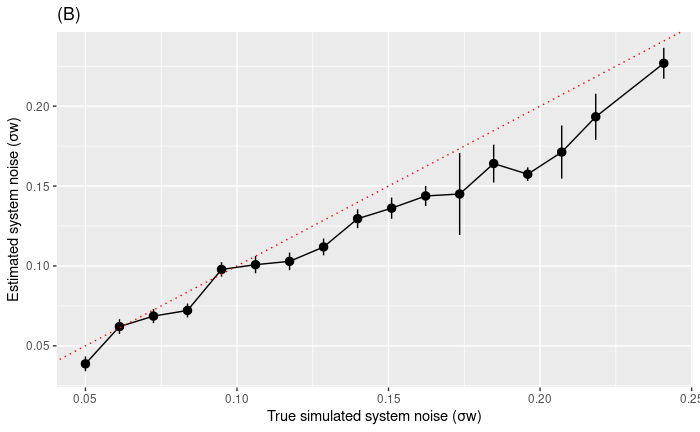

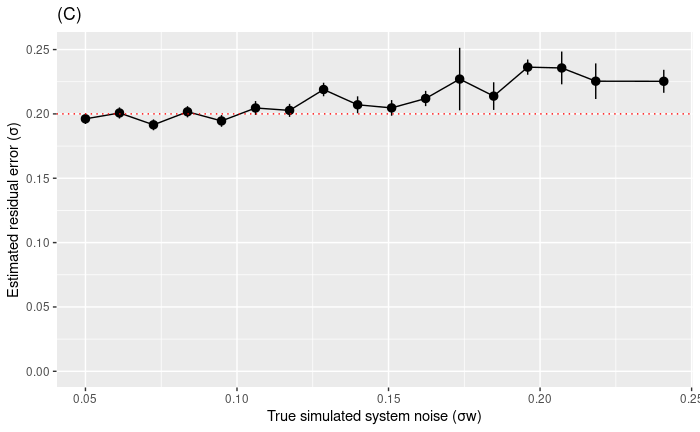

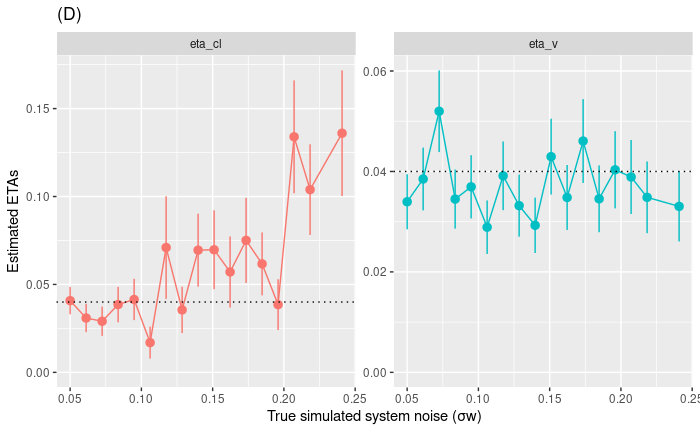

**Supplementary Code 1: NONMEM ODE Reference Model for One-Compartment IV PK Data**

$PROBLEM **NONMEM ODE Model**

$INPUT	ID TIME CONC=DV AMT MDV EVID CMT SDEe DVV

$DATA	fit.csv IGNORE=@

$SUB ADVAN1 TRANS2

$PK

MU_1 = THETA(1)

var1 = DEXP(MU_1 + ETA(1))

MU_2 = THETA(2)

var2 = DEXP(MU_2 + ETA(2))

V=var1

CL=var2

S1=V

$ERR

IPRE=(F)

Y=IPRE*(1+ERR(1))

IPRED=IPRE

$THETA

3.1 ; ; log(v)

1.1 ; ; log(cl)

$OMEGA

.03

.03

$SIG

.05

$EST METHOD=IMP INTERACTION NOABORT SIGL=5 PRINT=1 IACCEPT=1.0 CTYPE=3

$COV MATRIX=R UNCONDITIONAL

$TABLE ID TIME CONC AMT MDV EVID CMT SDEe DV IPRED

NOPRINT ONEHEADER FILE=xx1.csv

**Supplementary Code 2: NONMEM SDE Model for One-Compartment IV PK Data with System Noise Added to the Disposition Component**

$PROBLEM	NONMEM SDE Model for One-Compartment IV PK Data with System Noise Added to the Disposition Component

$ABBR PROTECT

$ABBR DES=FULL

$INPUT	ID TIME CONC=DV AMT MDV EVID CMT SDE DVV

$DATA	fit.csv IGNORE=@

$SUBROUTINE ADVAN13 TOL=9 ATOL=9

OTHER=/global/pkms2/data/Test/test.yan.li/00_yan_2023/06_02_2023_sde/sde.f90

$ABBR DECLARE SGW(1)

$MODEL

COMP=(central)

COMP=(OBSQ1)

COMP=(P1)

$PK

IF(NEWIND.NE.2) OT = 0

MU_1 = THETA(1)

var1 = DEXP(MU_1 + ETA(1))

MU_2 = THETA(2)

var2 = DEXP(MU_2 + ETA(2))

var3 = DEXP(THETA(3))

var4 = DEXP(THETA(4))

v=var1

cl=var2

kel=cl/v

sig=var3

SIGW1 = var4

NCMT = 1.0

NDE = 1.0

DEL=10**(-12)

$DES

FIRSTEM = 1

DADT(1) = -kel*A(1)

DADT(2)=A(1)/v ;DLOG((A(1)+DEL)/v)

SGW(1)= SIGW1 ; for additive and SIGW1*A(1) for proportional

"LAST

"	CALL SDE_DER(DADT,A,DA,IR,SGW,NDE,NCMT)

$ERROR (OBS ONLY)

IPRED = A(1)/v ;log((A(1)+DEL)/v)

W	= sig*IPRED

WS	= 1000.0D0

Y	= IPRED + W*EPS(1) + WS*EPS(2)

"LAST

"	CALL SDE_CADD(A,HH,TIME,DV,CMT,NDE,NCMT,SDE)

$THETA

3.1 ; ; log(v)

1.1 ; ; log(cl)

-1.5 ; ;(0, 0.3) ; log(sig)

0.95 ; ; (0, .02) ;log(sigw)

$OMEGA

.03

.03

$SIGMA

1 FIX

1 FIX

$EST METHOD=IMP INTERACTION NOABORT SIGL=5 PRINT=1 IACCEPT=1.0 CTYPE=3

$COV MATRIX=R UNCONDITIONAL

$TABLE ID TIME CONC AMT MDV EVID CMT SDE DV IPRED

NOPRINT ONEHEADER FILE=xx0.1.csv

**Supplementary Code 3: NONMEM SDE Model for One-Compartment IV PK Data with System Noise Added to the Elimination Process.**

$PROBLEM	NONMEM SDE Model for One-Compartment IV PK Data with System Noise Added to the Elimination Process

$ABBR PROTECT

$ABBR DES=FULL

$INPUT	ID TIME CONC=DV AMT MDV EVID CMT SDE DVV

$DATA	fit.csv IGNORE=@

$SUBROUTINE ADVAN13 TOL=9 ATOL=9

OTHER=/global/pkms2/data/Test/test.yan.li/00_yan_2023/06_02_2023_sde/sde.f90

$ABBR DECLARE SGW(2)

$MODEL

COMP=(central)

COMP=(logkel)

COMP=(OBSQ1)

COMP=(P1)

COMP=(P2)

COMP=(P3)

$PK

IF(NEWIND.NE.2) OT = 0

MU_1 = THETA(1)

var1 = DEXP(MU_1 + ETA(1))

MU_2 = THETA(2)

var2 = DEXP(MU_2 + ETA(2))

var3 = DEXP(THETA(3))

var4 = DEXP(THETA(4))

v=var1

cl=var2

kel=cl/v

sig=var3

SIGW1 = var4

NCMT = 1.0

NDE = 2.0

DEL=10**(-12)

A_0(2) = DLOG(kel)

$DES

FIRSTEM = 1

KELT = DEXP(A(2))

DADT(1) = -KELT*A(1)

DADT(2) = 0

DADT(3)=A(1)/v ;DLOG((A(1)+DEL)/v)

SGW(1)=0

SGW(2)= SIGW1

"LAST

"	CALL SDE_DER(DADT,A,DA,IR,SGW,NDE,NCMT)

$ERROR (OBS ONLY)

IPRED = A(1)/v ;log((A(1)+DEL)/v)

W	= sig*IPRED

WS	= 1000.0D0

Y	= IPRED + W*EPS(1) + WS*EPS(2)

A2=A(2)

"LAST

"	CALL SDE_CADD(A,HH,TIME,DV,CMT,NDE,NCMT,SDE)

$THETA

3.1 ; ; log(v)

1.1 ; ; log(cl)

-1.5 ; ;(0, 0.3) ; log(sig)

-2.0; logsigw_conc

$OMEGA

.03

.03

$SIGMA

1 FIX

1 FIX

$EST METHOD=IMP INTERACTION NOABORT SIGL=5 PRINT=1 IACCEPT=1.0 CTYPE=3

$COV MATRIX=R UNCONDITIONAL

$TABLE ID TIME CONC AMT MDV EVID CMT SDE DV A2 IPRED

NOPRINT ONEHEADER FILE=xx.1.csv

**Supplementary Code 4: NONMEM SDE Model for One-Compartment IV PK Data with System Noise Added to Both the Disposition Component and the Elimination Process.**

$PROBLEM	NONMEM SDE Model for One-Compartment IV PK Data with System Noise Added to Both the Disposition Component and the Elimination Process

$ABBR PROTECT

$ABBR DES=FULL

$INPUT	ID TIME CONC=DV AMT MDV EVID CMT SDE DVV

$DATA	fit.csv IGNORE=@

$SUBROUTINE ADVAN13 TOL=9 ATOL=9

OTHER=/global/pkms2/data/Test/test.yan.li/00_yan_2023/06_02_2023_sde/sde.f90

$ABBR DECLARE SGW(2)

$MODEL

COMP=(central)

COMP=(logkel)

COMP=(OBSQ1)

COMP=(P1)

COMP=(P2)

COMP=(P3)

$PK

IF(NEWIND.NE.2) OT = 0

MU_1 = THETA(1)

var1 = DEXP(MU_1 + ETA(1))

MU_2 = THETA(2)

var2 = DEXP(MU_2 + ETA(2))

var3 = DEXP(THETA(3))

var4 = DEXP(THETA(4))

var5 = DEXP(THETA(5))

v=var1

cl=var2

kel=cl/v

sig=var3

SIGW1 = var4

SIGW2 = var5

NCMT = 1.0

NDE = 2.0

DEL=10**(-12)

A_0(2) = DLOG(kel)

$DES

FIRSTEM = 1

KELT = DEXP(A(2))

DADT(1) = -KELT*A(1)

DADT(2) = 0

DADT(3)=A(1)/v ;DLOG((A(1)+DEL)/v)

SGW(1)= SIGW1 ; ; for additive and SIGW1*A(1) for proportional *(A(1)+DEL)

SGW(2)= SIGW2

"LAST

"	CALL SDE_DER(DADT,A,DA,IR,SGW,NDE,NCMT)

$ERROR (OBS ONLY)

IPRED = A(1)/v ;log((A(1)+DEL)/v)

W	= sig*IPRED

WS	= 1000.0D0

Y	= IPRED + W*EPS(1) + WS*EPS(2)

A2=A(2)

"LAST

"	CALL SDE_CADD(A,HH,TIME,DV,CMT,NDE,NCMT,SDE)

$THETA

3.1 ; ; log(v)

1.1 ; ; log(cl)

-1.5 ; ;(0, 0.3) ; log(sig)

-2.0; logsigw_conc

-3 ; ; (0, .02) ;log(sigw_kel)

$OMEGA

.03

.03

$SIGMA

1 FIX

1 FIX

$EST METHOD=IMP INTERACTION NOABORT SIGL=5 PRINT=1 IACCEPT=1.0 CTYPE=3

$COV MATRIX=R UNCONDITIONAL

$TABLE ID TIME CONC AMT MDV EVID CMT SDE DV A2 IPRED

NOPRINT ONEHEADER FILE=xx.1.csv
